# Metabolic responses to food and temperature in deep-sea isopods, *Bathynomus doederleini*

**DOI:** 10.1101/2023.01.23.525261

**Authors:** Shogo Tanaka, Yurika Ono, Shin-ichiro Tanimae, Toru Moriyama, Shingo Fujimoto, Mitsuharu Yagi

## Abstract

Metabolic rate, the energy required per unit of time for an organism to sustain life, is influenced by both intrinsic and extrinsic factors. Despite the similarities among living organisms across the various domains of life, it has been observed that those adapted to deep-sea environments exhibit notable distinctions from those in shallower waters, even when accounting for size and temperature. However, as deep-sea organisms are infrequently kept in captivity for prolonged periods, investigations into their potential metabolic responses to food and temperature have yet to be conducted. In this study, we demonstrate the impact of food (specific dynamic action: *SDA*) and temperature (*Q_10_*) on the metabolic rate of the deep-sea isopod *Bathynomus doederleini*. Positive correlations were found between *SDA* parameters (peak, time to peak, duration, and factorial scope) and meal size in deep-sea organisms. The postprandial metabolic rate, at a meal size of 45.4%, increased by approximately 6.5-fold, and the duration was 20 days. Within the temperature range of their natural habitat, the overall *Q_10_* was 2.36, indicating that a 10 °C increase would lead to a 2.4-fold increase in resting metabolic rate. The mean metabolic rate of this species, corrected for the equivalent temperature, was significantly 63% lower than the metabolic scaling rule for aquatic invertebrates. This low metabolic rate suggests that deep-sea isopods can survive for a year on a mere few grams of whale blubber at a water temperature of 10.5 °C. This information is crucial for understanding the metabolic strategies and consequences of adaptation to a deep-sea environment.

## 1. Introduction

Aerobic organisms sustain themselves by utilizing oxygen to metabolize nutrients in their bodies and generate energy. The metabolic rate, which is the energy expenditure per unit time required for an organism to survive, is often referred to as the “fire of life.” Energy metabolism is largely constrained by physical and kinetic factors, such as body size and temperature (Peters, 1983; Calder, 1984; Schmidt-Nielsen, 1984; Blaxter, 1989; Brown and West, 2000; Glazier, 2005). The rate of increase in metabolic rate for a 10 °C increase in temperature is known as the *Q_10_* (Schmidt-Nielsen, 1997). It has been posited that metabolic rate may also be influenced by factors such as an organism’s activity level (Biro and Stamps, 2010), ontogenetic stage (Yagi et al., 2010; Yagi and Oikawa, 2014), taxonomic and phylogenetic status (Hayssen and Lacy, 1985), geographic range and distribution patterns (Lovegrove, 2000), habitat temperature conditions (MacMillen and Garland, 1989), and the availability and preferences of resources (McNab, 1986).

The correlation between resting metabolic rate and body size after adjusting for temperature, known as metabolic scaling, has been the subject of extensive research for a considerable period. In his seminal work, Hemmingsen (1960) established that the following allometric equation could adequately express the relationship between metabolic rate and body size across various taxa:

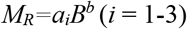

where MR represents resting metabolic rate, B denotes body mass, a represents a scaling constant and b signifies the scaling exponent. The value of *i* denotes the three different animal groups (homeotherms, heterotherms and unicellular organisms), which exhibit distinct values for the scaling constant (intercepts). Despite ongoing debate regarding the scaling exponent (Kozłowski and Konarzewski, 2004; O’Connor et al., 2007; Harrison, 2017; Glazier, 2022), it has been established that each of these animal groups displays a consistent trend (Makarieva et al., 2008).

Is the “fire of life” less ardent in deep-sea organisms? The deep sea environment is characterized by decreasing temperatures and increasing pressures with increasing water depth, as well as being oligotrophic and having a significantly reduced light intensity (Merrett and Haedrich, 1997). Some deep-sea organisms that have adapted to such conditions exhibit lower metabolic rates than their shallow-water counterparts, even after accounting for temperature and body size adjustments (Seibel and Drazen, 2007). The “visual-interaction hypothesis” has been proposed to explain this phenomenon (Childress and Mickel, 1985; Seibel and Drazen, 2007). This hypothesis posits that visually mediated predation and prey behaviour diminish with increasing depth (reduced light), rendering active swimming redundant in deeper and darker environments. As a result, metabolic activity has evolved to decrease with depth in visually developed organisms (Seibel and Drazen, 2007). This trend has been observed in cephalopods, crustaceans with well-developed vision, and teleost fish (Sullivan and Somero, 1980; Seibel et al., 1997; Seibel and Drazen, 2007).

Even among deep-sea organisms, metabolic reductions may not have evolved in the deep-sea isopod *Bathynomus doederleini*. This species, which possesses an adult size of approximately 100 mm, is primarily found in the Pacific Ocean, ranging from Japan to the Philippines at depths of 200 - 600m (Sekiguchi et al., 1981). Deep-sea isopods are considered benthic scavengers and typically burrow in the seafloor to evade predation (Matui et al., 2011). The “visual-interaction hypothesis” suggests that animals with poor visual acuity and those inhabiting the benthos do not exhibit metabolic reductions with increasing depth (Seibel and Drazen, 2007). Indeed, it has been reported that metabolic rate does not decrease with increasing depth in benthic crustaceans, octopods, chaetognaths, medusa, and worms, whose behaviour is not as reliant on vision (Thuesen and Childress, 1993a; Thuesen and Childress, 1993b; Seibel and Childress, 2000; Seibel and Drazen, 2007). However, to the best of our knowledge, there are no reported metabolic rates for this deep-sea species.

Determining the metabolic rate of deep-sea organisms in response to environmental changes presents a formidable challenge, as sudden fluctuations in water pressure can prove fatal during the process of capturing and raising them to the surface. Despite the existence of reports of *in vitro* enzyme activity (Childress and Somero, 1979) and measurements under high-pressure conditions (Mickel and Childress, 1982) pertaining to deep-sea metabolism, the response of these organisms to varying environmental conditions remains inadequately understood. Conversely, some studies have conducted *in situ* measurements of metabolism in deep-sea environments, yielding intriguing insights (e.g. Bailey et al., 2002). However, metabolic rates that take into account the effects of specific dynamic actions (*SDA*) have yet to be determined, as the feeding state of the individuals being measured cannot be controlled. *SDA* refers to the increase in metabolic rate following feeding (McCue, 2006). In order to exclude the effects of *SDA*, resting metabolic rate should be measured when individuals are fasting (Kleiber, 1932; Wang et al., 2006; Secor 2009). Notably, no studies on *SDA* in deep-sea organisms have been conducted to date.

Deep-sea isopods possess a durable exoskeleton and lack a swim bladder, rendering them tolerant to fluctuations in water pressure, indicating their ability to persevere for extended intervals under ambient pressure, thus facilitating thorough measurements to be conducted under varying conditions. The overarching objectives of the current study were to investigate: (i) the impact of food intake (*SDA*) and (ii) the effect of water temperature (*Q_10_*) on the metabolic rate of the deep-sea isopod. This knowledge is crucial for comprehending the physiology of deep-sea organisms, specifically metabolic strategies pertaining to adaptation to the deep-sea milieu, and ecological and evolutionary bioenergetics.

## 2. Materials and methods

### 2.1. Ethics statement

All experiments were performed in conformity with Japanese Animal Care protocols and received approval from the Nagasaki University Fish and Invertebrate Experimental Ethics Committee (Ethics Approval No. NF-0069).

### 2.2. Animals

Deep-sea isopods were obtained through the efforts of local fishermen utilizing bait traps in Suruga Bay and off the coast of Goto Island, Japan in 2021. A total of forty-six specimens (38 from Suruga Bay and 8 from Goto Island) were conveyed in refrigerated storage to the Fish and Ships Lab, Faculty of Fisheries at Nagasaki University. Upon arrival, they were promptly transferred to a 200 L black circular tank (diameter: 0.8 m) until the initiation of experiments. The rearing tank was equipped with aeration, a filtration system (Ehim 2260, Ehime, Germany), and cooled via a cooling chiller (AZ280X, Rei-Sea, Tokyo, Japan). The deep-sea isopods were provided with sustenance in the form of swordtip squid *Uroteuthis edulis* once every two weeks, with the exception of the experimental period. The rearing water was replaced by half every seven days. The rearing room was consistently shaded, and observations and experimental settings were conducted under red light. Temperature Data Loggers (model UTBi-001, Onset Computer Corporation, MA, U.S.A.) were submerged in both the rearing and experimental tanks. During the rearing period, the mean water temperature (± S.D.) was 12.1 ± 0.2 °C and salinity ranged from 33.1 - 34.3 PSU.

### 2.3. Respirometory

Oxygen consumption during resting status of animals was measured as a proxy of resting metabolic rate (Yagi et al., 2010). The technique of intermittent-flow respirometry as described by Mochnac et al. (2017) was employed to measure oxygen consumption. The respirometry unit, with a volume of approximately 80 L, consisted of a respiration chamber and a water bath, the temperature of which was regulated by means of a heater and cooler, and was filled with air-saturated seawater. The respiration chamber was cylindrical in shape (90 mm × 200 mm, 930 mL), fabricated from acrylic and stirred by a magnetic stirrer. Dissolved oxygen and salinity were measured using an optical multimeter (Multi3430, Weilheim, Germany). Measurements were taken at intervals of 50-85 minutes (with close and open times of 20-45 minutes and 30-40 minutes, respectively). The dissolved oxygen never dropped below 80%. In parallel, a blank control was conducted to calculate the background respiration (which never exceeded 3%). The respiration chamber was covered with a blackout curtain during measurements.

*MO_2_* (expressed in units of mgO_2_ L^−1^ min^−1^) was computed by taking into account the decrease in dissolved oxygen in both the experimental chamber and the blank (absent of animals) as follows:

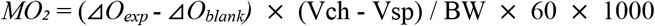

where ⊿*O*_exp_ and ⊿*O*_blank_ (mgO_2_ L^−1^ min^−1^) are the gradients of linear regression of dissolved oxygen in the experimental and blank respirometry in relation to the duration of incubation. *V_ch_* denotes the volume of the respiration chamber (930 mL), and *V_sp_* represents the volume of the specimen, determined from its body weight. The density of the deep-sea isopods was found to be 1.225 g mL^−1^ as determined by the volumetric method utilising Archimedes’ principle.

### 2.4. Effects of food (SDA)

To examine the effects of food on metabolic rate (*SDA*), pre- and post-feeding oxygen consumption were assessed. The measurements were conducted at a temperature of 12.1 ± 0.3 °C for a period of 24 hours prior to feeding using individuals that fasted for more than 28 days. The mean wet body weight (± S.D.) and body length were 33.0 ± 4.6 g and 100.0 ± 3.8 mm (n = 14), respectively. The food item utilized was swordtip squid. In order to achieve significant variation in feeding rate, the feeding time ranged from 2 to 10 minutes, resulting in a proportion of 6.8 - 45.4% of the body weight. The weight of the feed was computed by subtracting the weight of the food prior to and after feeding. The period of measurements varied from 6 to 30 days depending on food intake. During the measurement period, 10 L of seawater in the respiratory tank was changed daily without altering the water temperature.

### 2.5. Effects of temperature (Q_10_)

To investigate the effects of temperature on metabolic rate (*Q_10_*), oxygen consumption was measured at four distinct temperatures (15, 12, 9 and 6 °C) which are within the temperature range in which the species is found in the natural habitat (Yagi et al., unpublished data). Five individuals were randomly selected from the bloodstock tank and placed in three separate 30 L black circular tanks (diameter: 0.4 m) that were independent of one another. Each tank was equipped with aeration, a filtration system (Eheim Classic Filter 2213, Deizisau, Germany), and a cooler (Cool Way BK110, Gex, Osaka, Japan). Ten L of seawater was exchanged every week without altering the temperature in each tank. Prior to the metabolic rate measurements, the individuals were acclimatized for a minimum of 14 days at each temperature. The measurements were carried out in descending order from high (15 °C) to low (6 °C) temperatures, with the temperatures being decreased to 1 °C per day. During each acclimation period, the mean water temperature (± S.D.) was 15.3 ± 0.2 °C, 12.2 ± 0.2 °C, 9.2 ± 0.2 °C, 6.2 ± 0.2 °C. The aforementioned experiment was repeated twice in different populations.

Measurements were performed on a total of 25 individuals at temperatures of 15 °C (n = 8), 12 °C (n = 10), 9 °C (n = 8), and 6 °C (n = 9). The mean wet body weight (± S.D.) and body length (± S.D.) were 34.0 ± 2.9 g and 102.7 ± 5.6 mm (n = 25), respectively. Individuals underwent a fast of at least 14 days prior to the measurement. Prior to the measurements, individuals were acclimated to the respiration chamber overnight and oxygen consumption was measured three times in each individual.

### 2.5. Data analysis

The variables commonly employed to quantify *SDA* - were calculated according to Secor (2009) and Feher (2017), utilizing the following model equations:

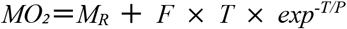

Where *MO_2_* (mgO_2_ kg^−1^ h^−1^) represents metabolic rate, *M_R_* denotes the resting *MO_2_* averaged over the 6 hours preceding feeding, *F* represents meal size (the proportion (%) of food intake (g) divided by the individual body weight (g)), *T* (h) denotes the time elapsed since feeding, and *P* (h) represents the time until peak. Thus, the maximum of the curve occurs at *T* = *P* at an increment of *F T exp*^−1^ (Feher, 2017). *MO_2_* data between 05:00 AM and 17:00 PM were analyzed in accordance with Roe (2004). Subsequently, the four parameters were determined (Fig. 1), i.e., (i) peak; postprandial peak in metabolic rate, (ii) time to peak; duration from time of feeding to peak metabolic rate, (iii) duration; time from feeding when metabolic rate achieves a 5% increase in resting metabolic rate, (iv) factorial scope; postprandial peak divided by resting metabolic rate. Linear relationships between *SDA* parameters and meal size were analyzed by Pearson’s correlation coefficient test.

**Fig. 1.**
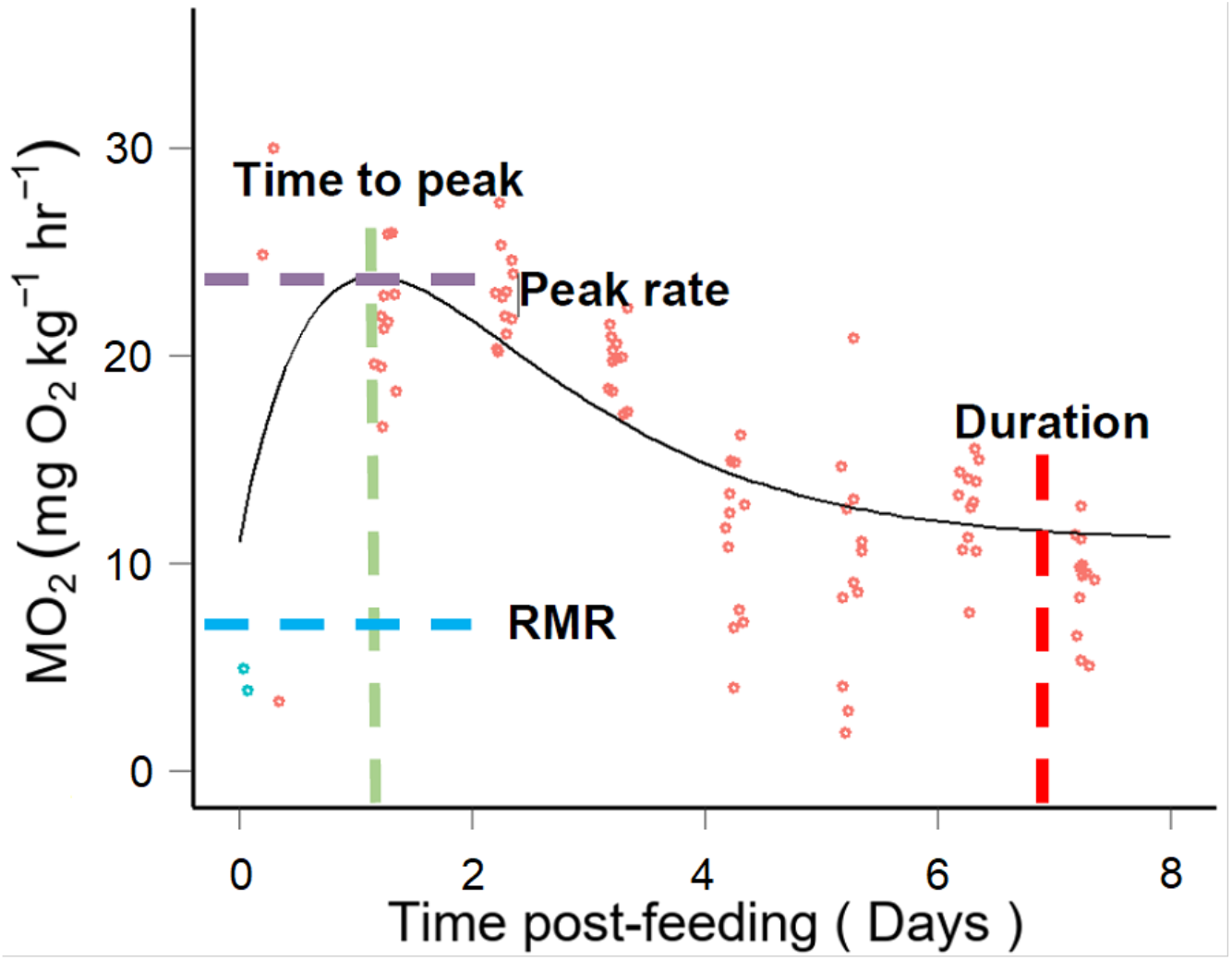
Impact of food intake on the metabolic rate of the deep-sea isopod *Bathynomus doederleini*. An illustration of the postprandial metabolic profile of metabolic rate plotted against time post-feeding (meal size was 11.5 % of body weight). Quantified specific dynamic action (*SDA*) parameters are highlighted.

An exponential curve was fitted to the effects of temperature on metabolic rate, with metabolic rate as the dependent variable and temperature as the independent variable. *Q_10_* was calculated by utilizing the following equation:

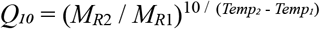

where *M_R1_* and *M_R2_* are the known metabolic rates (mgO_2_ kg^−1^ h^−1^) at their respective temperatures, *Temp_1_* and *Temp_2_* (°C) (Schmidt-Nielsen, 1997).

To verify the normality and homogeneity of the dataset for the application of parametric analysis, Shapiro-Wilk and Bartlett tests were conducted. All statistical analyses were performed using R (Version 4.2.2).

## 3. Results

### 3.1 Effects of food (SDA)

Postprandial metabolic rates of deep-sea isopods increased (Fig. 1). Mean resting metabolic rate (*M_R_* ± S.D.) was 9.5 ± 5.6 mgO_2_ kg^−1^ h^−1^, ranging from 3.9 to 21.6 mgO_2_ kg^−1^ h^−1^ (n = 14).

All *SDA* parameters displayed a dependency on meal size, with statistical significance (peak; t = 4.39, p < 0.001, time to peak; t = 2.19, p = 0.049, duration; t = 2.53, p = 0.026, factorial scope; t = 2.75, P = 0.018) (Fig. 2). Peak and time to peak ranged from 18.6 to 51.6 (Fig. 2a), and 0.5 to 5.6 (Fig. 2b), respectively. Duration and factorial scope ranged from 3.1 to 33.3 (Fig. 2c), and from 0.9 to 12.0 (Fig. 2d), respectively.

**Fig. 2.**
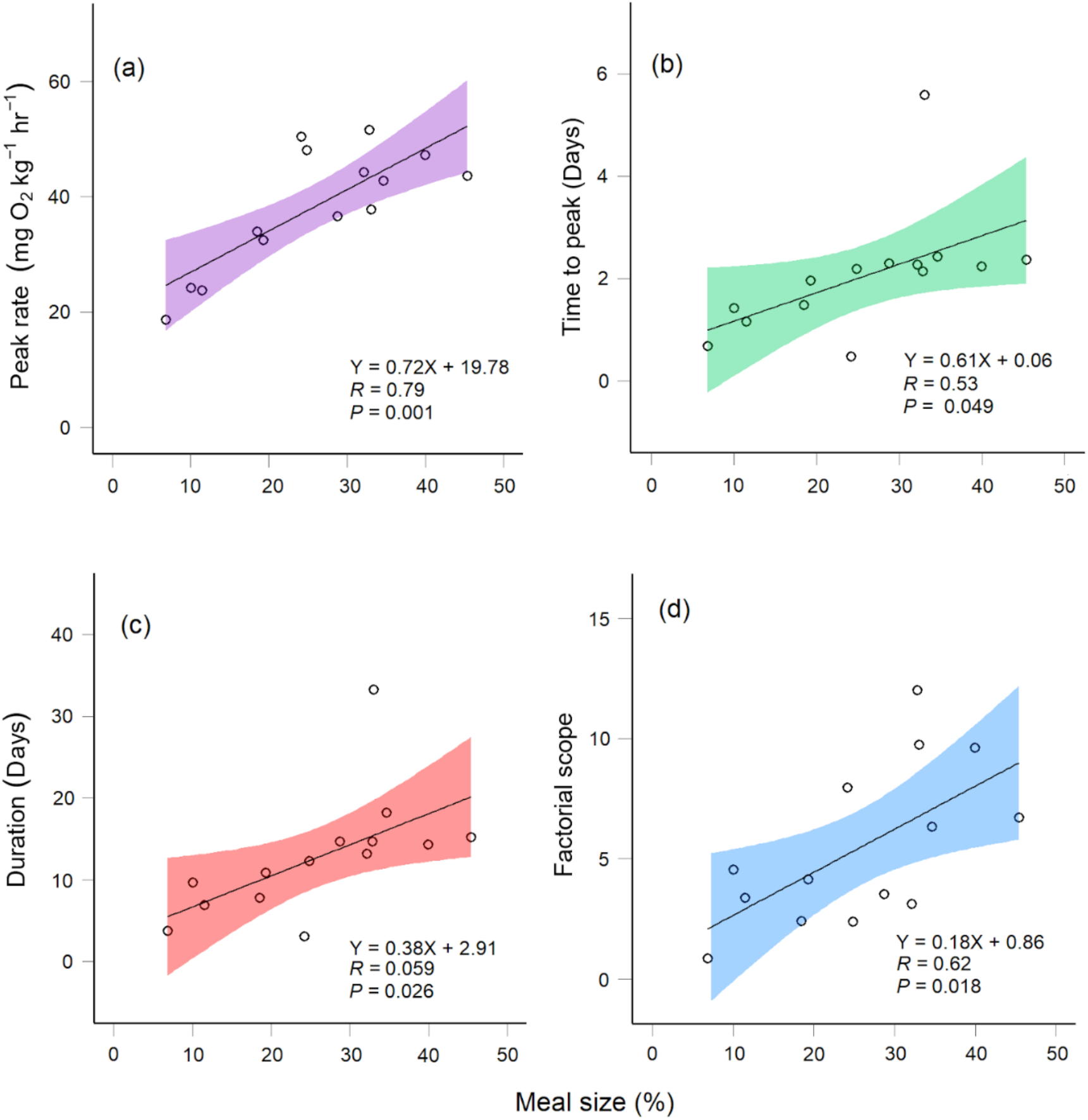
Relationships between specific dynamic action (*SDA*) parameters and meal size for deep-sea isopods *Bathynomus doederleini*. (a) Peak. (b) Time to peak. (c) Duration. (d) Factorial scope. The shaded areas indicate the 95% confidence interval for the fitted exponential curve.

### 3.2. Effects of temperature (Q_10_)

The resting metabolic rate significantly increased with increasing temperature (Fig. 3). The mean resting metabolic rate (± S.D.) at each temperature was 20.2 ± 4.1 (15 °C, n = 8), 15.8 ± 4.9 (12 °C, n = 10), 12.6 ± 5.5 (9 °C, n = 8) and 9.1 ± 2.9 mgO_2_ kg^−1^ h^−1^ (6 °C, n = 9). The overall *Q_10_* was 2.36, and the one at each temperature range was 2.20 (15-12 °C), 1.80 (12-9 °C) and 3.30 (9-6 °C).

**Fig. 3.**
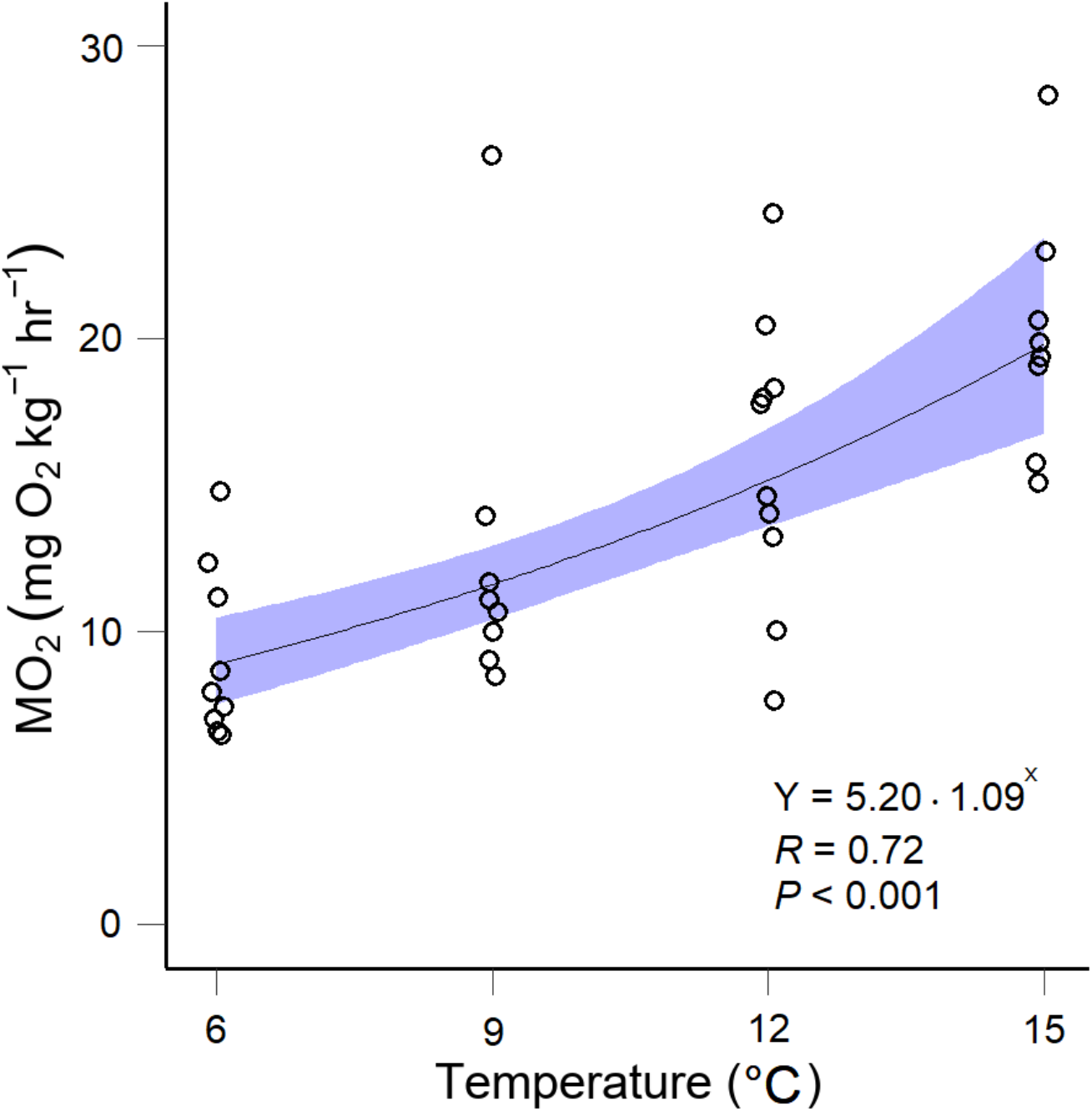
Effects of temperature on resting metabolic rate (*MO_2_*) in deep-sea isopods *Bathynomus doederleini*. The overall *Q_10_* was 2.36. The shaded area indicates the 95% confidence interval for the fitted exponential curve.

## 4. Discussion

This study quantified the responses of food and temperature on metabolic rate in deep-sea isopods through long-term rearing and temperature acclimation experiments. To the best of our knowledge, this is the first report of a positive correlation between *SDA* parameters (peak, time to peak, duration, and factorial scope) and meal size in deep-sea organisms. The postprandial metabolic rate at a meal size of 45.4% increased by approximately 6.5-fold and the duration was 20 days. Within the water temperature range of the species’ natural habitat, the overall *Q_10_* was 2.36, indicating that a 10 °C increase would result in a 2.36-fold increase in resting metabolic rate.

Deep-sea organisms may possess *SDA* characteristics that reflect their feeding strategy of consuming large quantities of food infrequently, due to the scarcity of resources in their deep-sea habitat. The postprandial increase in metabolism, known as *SDA*, reflects the energy requirements associated with digestion, including the transport, absorption, and storage of nutrients, as well as increased synthesis of proteins and lipids (Secor, 2009). This study confirms that deep-sea isopods also possess *SDA*, similar to shallow-water organisms. The duration of *SDA* increased with increasing meal size (Chakrabborty et al.,1992; Janes and Chappell, 1995; Toledo et al., 2003), which is consistent with the findings of this study. However, the calculated duration of this species was notably long, reaching up to 33.3 days (Fig. 2). This may reflect the substantial difference in meal size (33.0%), which is an order of magnitude greater than that of shallow-water organisms. A satiating meal typically ranges between 2 - 4% of an organism’s body weight (Elner, 1980; Robertson et al., 2002), and a 5% meal size would be considered high for crustaceans (Curtis et al., 2010). For comparison, the American crayfish *Procambarus clarkii* and the shallow-water crab *Hemigrapsus nudus*, which have similar body sizes to the deep-sea isopod, have been reported to have maximum meal sizes of 0.5% and 3.0%, respectively, with durations of 0.47 and 2.51 days (McGaw and Curtis, 2013). Furthermore, the deep-sea isopods in this study consumed a meal size of 45.4% within 10 minutes, indicating a high feeding rate. These adaptations allow deep-sea isopods to cope with the intense competition for patchy resources by rapidly consuming as much food as possible to avoid predation or cannibalism (Barradas-Ortiz et al., 2003; Smith and Baldwin, 1982).

Due to their well-developed digestive systems, deep-sea isopods may adopt an energetic strategy of increasing the factorial scope for a single large meal size. In this study, it was observed that the factorial scope of this species increased significantly with increasing meal size (Fig. 2d). Many organisms, including isopods, have been found to have a factorial scope within the range of 1.5 - 3.0 (Secor, 2009). However, this deep-sea isopod species exhibited a maximum factorial scope of 12.1. The presence of three pairs of hepatopancreatic glands in the oesophagus, which secrete digestive enzymes, and the likely use of energy-requiring absorptive processes, facilitated by a well-developed circulatory system (Kihara and Kuwasawa, 1984), likely contribute to the deep-sea isopod’s ability to consume large meals. However, this increased energy expenditure for *SDA* may also constrain behavioural activity. Indeed, post-feeding individuals of many meal sizes were observed to be rounded and static. McCue (2006) suggested that *SDA* scope can be compared to maximal metabolic scope to estimate the residual capacity for activity during digestion. Interestingly, it is plausible that there exists a trade-off relationship where more energy is allocated towards *SDA*, and behavioural activity is constrained.

Although deep-sea isopods inhabit deep-oceanic environments, this species exhibits a degree of adaptability to a relatively broad range of temperatures. The *Q_10_* coefficient, which measures the sensitivity of an organism’s metabolic rate to temperature fluctuations, has been observed to increase in value as temperatures approach their lethal limit (Christensen et al., 2021). The *Q_10_* of deep-sea isopods in this study was found to be 2.36 within the temperature range of 6 - 15 °C, a value comparable to that of many other organisms, typically falling within the range of 2 - 3 (Schmidt-Nielsen, 1997). However, no significant deviation in *Q_10_* value was observed at temperatures approaching the lethal limit, indicating that deep-sea isopods possess a high degree of adaptability to these temperature ranges. Indeed, deep-sea isopods are primarily found at depths of 200 - 600 meters (Sekiguchi et al., 1981). The water temperature immediately above the seafloor in the East China Sea, where this species occurs, averages 13.8 °C at a depth of 200 meters and 6.6 °C at 600 meters (Yagi et al., unpublished). This study provides further support for the adaptive nature of this species to the coastal-bathyal environment, particularly in terms of thermo-metabolic physiology.

The fire of life in deep-sea isopods was faint. The visual interaction hypothesis posits that the metabolic rate of swimming crustaceans and cephalopods significantly diminishes with increasing depth of habitat, in contrast to a lack of decline in benthic crustaceans and polychaetes (Seibel and Drazen, 2007). Given that this particular species is benthic, it is predicted that its metabolic rate should not deviate from that of shallow-water animals, according to the hypothesis. However, the mean metabolic rate of this species, adjusted for temperature (25 °C), was significantly 63% lower than the established metabolic scaling relationship for the aquatic invertebrate group (Makarieva et al., 2008) (Fig. 4). Decreases in metabolic rate have also been documented in benthic carrideans and crustaceans with increasing depth (Company and Sarda, 1998; Seibel and Drazen, 2007), in conformity with the present study. At present, additional information on metabolic rate and lifestyle is necessary to validate the hypothesis (Drazen and Seibel, 2007).

**Fig. 4.**
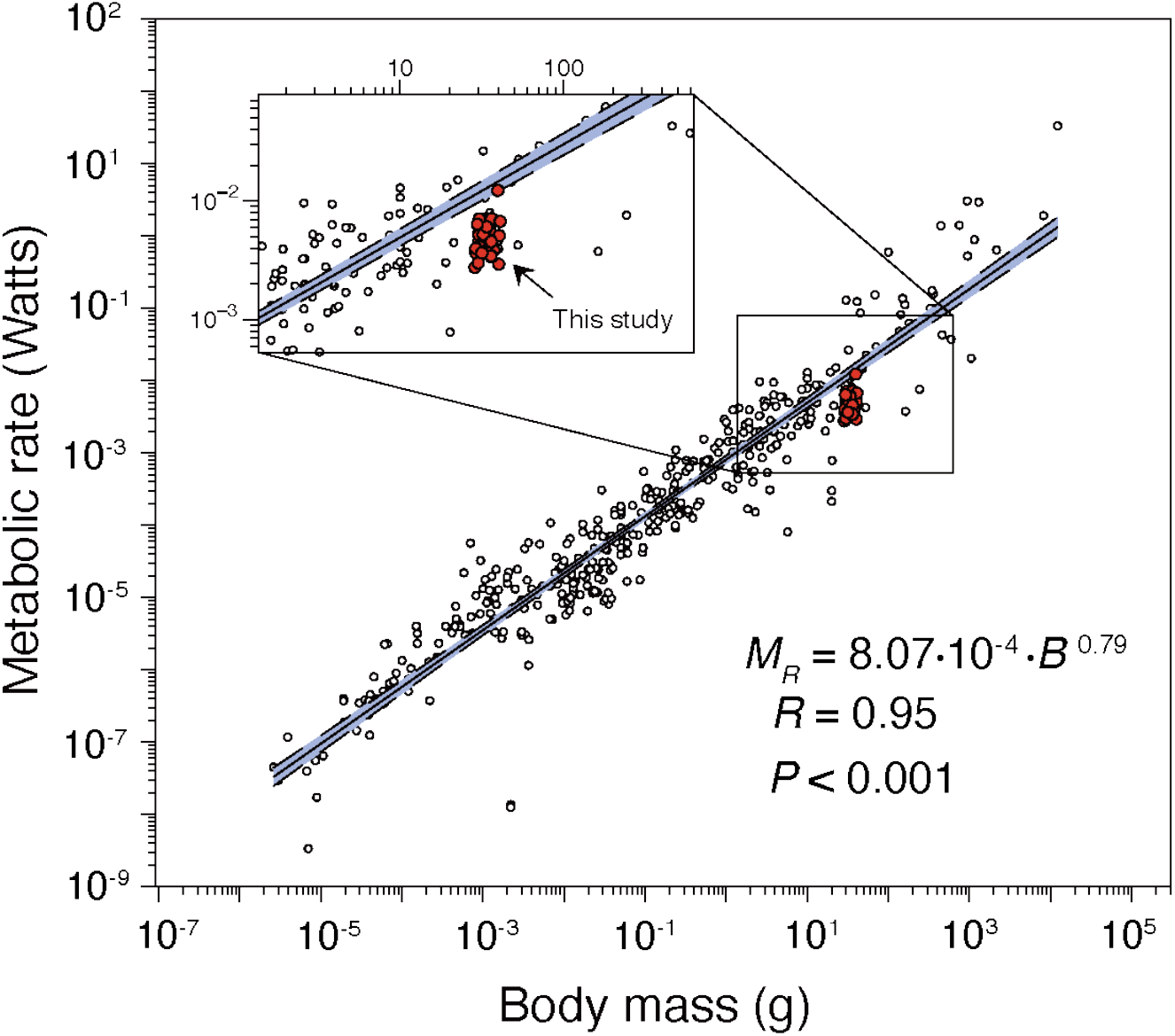
Metabolic scaling (the relationship between metabolic rate and wet body mass) in the deep-sea isopod *Bathynomus doederleini* in comparison to published interspecific comparisons (Makarieva et al., 2008) amongst aquatic invertebrates. Measurements from aquatic invertebrates (n = 376) and our measurements of deep-sea isopod (n = 35; represented by red circles) were adjusted to 25 °C, using the factor *Q_10_* = 2, in accordance with Makarieva et al. (2008). The regression line and its 95% confidence interval (shaded area) did not encompass the data from this study.

According to calculations based on the low metabolic rate of the deep-sea isopod, this species may survive for a year on 2.1 grams of whale blubber at a temperature of approximately 10 °C. Given the species’ remarkable meal size (Fig. 2), it can be estimated that it would subsist for 6.6 years on a single diet, although the energy costs associated with foraging, growth, and reproduction would be an additional factor in reality. Indeed, Ginn et al. (2014) reported that the closely related giant isopod *B. giganteus* lived for five years in aquarium captivity, even after fasting. The deep-sea, low-metabolism colosal squid *Mesonychoteuthis hamiltoni* (with a body weight of 500 kg) has been estimated to consume only 30 grams of fish per day, regardless of its giant body size (Rosa et al., 2010). Nonetheless, the mechanisms underlying low metabolic rates in deep-sea organisms, including this species, remain poorly understood. It is suggested that mechanisms that preserve low metabolic rates that do not conform to metabolic scaling may exist, and further research into the metabolism of deep-sea organisms is crucial.

## Acknowledgements

This work was supported by JSPS KAKENHI (Grant Number JP18K14790 and JP21K06337 to M.Y.). We are grateful to the “Fish and Ships Laboratory” students from the Faculty of Fisheries, Nagasaki University, who assisted us with the study, and Hisashi Hasegawa, Kazutaka Hasegawa of the Chokane Maru, which belongs to Kogawa Port in Yaizu City, Shizuoka Prefecture, for capturing the deep-sea isopods. Finally, thanks to the editors and anonymous reviewers for their valuable comments and suggestions that greatly improved the quality of the manuscript.

